# Modulation of post-traumatic immune response using anti-IL-1 therapy for improved visual outcomes: Anti-IL-1 improves post-traumatic visual outcomes

**DOI:** 10.1101/659581

**Authors:** Lucy P. Evans, Addison W. Woll, Shu Wu, Brittany P. Todd, Nicole Hehr, Adam Hedberg-Buenz, Michael G. Anderson, Elizabeth A. Newell, Polly J. Ferguson, Vinit B. Mahajan, Mathew M. Harper, Alexander G. Bassuk

## Abstract

The purpose of this study was to characterize acute changes in inflammatory pathways in the mouse eye following a blast-mediated TBI (bTBI) model, and to determine if modulation of these pathways could protect the structure and function of retinal ganglion cells (RGC). bTBI was induced in C57BL/6J male mice by exposure to three 20 PSI blast waves, with an inter-blast interval of one hour. Acute cytokine expression in retinal tissue was measured through real-time quantitative polymerase chain reaction (RT-qPCR) 4 hours post-blast. Increased retinal expression of *lL-1β, IL-1α, IL-6*, and *TNFα* was observed in bTBI mice exposed to blast when compared to shams, which was associated with activation of microglia and macroglia reactivity, assessed via immunohistochemistry with IBA-1 and GFAP, respectively, 1 week post-blast. Inhibition of the IL-1 pathway was accomplished using anakina, an IL-1RI antagonist. Retinal function and structure were evaluated 4 weeks post-injury using pattern electroretinogram (PERG) and optical coherence tomography (OCT), respectively. After bTBI, anakinra treatment resulted in a preservation of RGC function and RGC structure when compared to saline treated bTBI mice. Optic nerve integrity analysis demonstrated a tred of decreased suggesting that IL-1 blockade also prevents axonal damage after blast. Blast exposure results in increased retinal inflammation including upregulation of pro-inflammatory cytokines and activation of resident microglia and macroglia. This may partially explain the RGC loss we observed in this model as blockade of the acute inflammatory response after injury with the IL-1R1 antagonist anakinra resulted in preservation of RGC function and structure.

**Significance Statement:** Blast-mediated traumatic brain injury (bTBI) affects military members and civilians as a direct result of combat, workplace accidents, or intentional terrorist attacks. The retina is a central nervous system (CNS) tissue that is vulnerable to blast exposure. Individuals with bTBI often report visual dysfunction, but the mechanisms of ocular injury are poorly understood. This study demonstrates that damaging neuroinflammation contributes to retinal injury following blast-mediated TBI. We also identified anakinra, an anti-IL-1 therapy currently utilized for other diseases, as a potential pharmacologic agent that could prevent ocular damage after blast. These findings will aide in the development of novel treatments for vision preservation.

## Introduction

Traumatic brain injury (TBI) is a leading cause of death and disability worldwide, which results in enormous social and economic costs. Due to the use of improvised explosive devices in 21^st^ century military conflicts, the number of blast-related injuries causing TBI (bTBI) has increased dramatically in both military personnel and civilians, while blast-related deaths have decreased due to enhanced protective equipment (Cockerham, Goodrich et al. 2009, Cockerham, Rice et al. 2011). Unfortunately, there are no effective pharmacologic therapies to prevent neuronal loss after blast, and treatment is limited to supportive care.

The retina is a central nervous system (CNS) tissue vulnerable to injuries affecting the brain (London, Benhar et al. 2013). This is particularly true in the setting of bTBI as the eye is also exposed to the primary blast wave. Many military personnel and civilians who suffer from bTBI also report symptoms of visual dysfunction with retinal pathology, which can present either acutely or chronically after the initial injury (Cho, Bakken et al. 2009), (Cockerham, Goodrich et al. 2009). Although bTBI patients report a wide range of visual disturbances, little is known about the molecular mechanisms driving visual dysfunction. Murine bTBI models display many of the long-term visual deficits observed in patients, particularly retinal ganglion cell (RGC) dysfunction and subsequent cell death (Mohan, Kecova et al. 2013, Dutca, Stasheff et al. 2014, Bricker-Anthony, Hines-Beard et al. 2016, Yin, Voorhees et al. 2016, Evans, Newell et al. 2018).

Following a TBI, secondary signaling cascades occur in the brain, including robust neuroinflammation (Simon, McGeachy et al. 2017), which can trigger progressive neuronal dysfunction, neuronal death, and tissue destruction that exacerbates the initial injury. A therapeutic window exists in the hours to days after initial trauma when inflammatory pathways are initiated and can be blocked to halt progressive tissue injury. Components implicated in this neuroinflammation include rapid upregulation of the IL-1 cytokine family, including IL-1α and IL-1β (Griffin, Sheng et al. 1994, Fan, Young et al. 1995, Hutchinson, O’Connell et al. 2007, Utagawa, Truettner et al. 2008), IL-1 cytokines can be produced via activated resident macroglia (astrocytes and Müller glia) and microglia, or through infiltration of systemic immune cells into the CNS (Allan, Tyrrell et al. 2005). The primary receptor for IL-1, IL-1RI, is present on many cell types, including both CNS and peripheral immune cells, which in turn may contribute to further neuronal injury.

Several preclinical studies have begun to test blockade of IL-1β following TBI, including use of anti-IL-1β antibody or blockade of IL-1RI. Encouraging results showed both histologic and functional improvement with decreased tissue loss and attenuated cognitive deficits (Toulmond and Rothwell 1995, Tehranian, Andell-Jonsson et al. 2002, Clausen, Hanell et al. 2009, Newell, Todd et al. 2018). Anakinra, a recombinant human interleukin-1 receptor antagonist (rhIL-1Ra) that mimics the action of the natural antagonist IL-1Ra, is also being studied as a potential therapy for TBI. In a phase-2 trial of adults with severe TBI, anakinra had a good safety profile and brain penetration when administered peripherally (Helmy, Guilfoyle et al. 2014).

The subsequent questions that remain are: what are the targets, what is the therapeutic window for effective intervention, and is it possible to pharmacologically treat bTBI patients to prevent long-term visual dysfunction? In this study we developed a model of repeated blast injury that did not initially show clinical symptoms with normal activity and demeanor— mimicking a very common human TBI scenario. We then identified molecular mediators of inflammation in the retina, which have long been implicated in inflammatory damage in the brain after TBI, and confirmed that retinal injury could be detected non-invasively using clinical testing modalities used in humans. Finally, we rescued this damage using a pharmacologic agent blocking the IL-1 receptor (IL-1RI), an effect that could be monitored by retinal exam. Taken together, this points to a diagnostic and therapeutic strategy for blast-induced retinal injury.

## Materials and Methods

### Animals

All animal studies were conducted in accordance with the ARVO Statement for the Use of Animals in Ophthalmic and Vision Research and were approved by the University of Iowa and Iowa City Veterans Affairs Instiutitonal Animal Care and Use committee. Studies were conducted on male C57BL/6J mice purchased from The Jackson Laboratory (Bar Harbor, ME) aged 2-4 months, with an average weight of 26.4 +/-1.3 g. A total of 125 mice were used for this study. Mice were housed under a 12-h light-dark cycle with *ad libitum* access to food and water.

### Blast Injury Induction

An enclosed blast chamber was used, one half of which was pressurized, as previously described (Mohan, Kecova et al. 2013). A plastic mylar membrane (Mylar A, 0.00142 guage; country Plastics, Ames, IA) was placed over a 13-cm opening that separates the sides of the chamber. The unpressurized side of the tank contained a padded polyvinyl chloride (PVC) protective restraint in which to place an anesthetized mouse 30 cm from the mylar membrane. Compressed air was pumped into the pressurized side of the chamber until the membrane ruptured at 20 ± 0.2 psi (137.8 ± 1.3 kPa, mean ± SEM), creating a blast wave. Human and murine response to TBI can be heterogeneous, so we administered three injuries to each mouse in order to decrease the variability of the inflammatory response within the retina. Mice were oriented within the chamber with the left side of the head positioned towards the blast wave (direct exposure) and the right side facing away from the direction of the blast wave (indirect exposure) and then exposed to three blast injuries, each one hour apart (**Fig. 1)**. The mouse’s body was shielded via the PVC restraint to limit blast wave pressure exposure primarily to the head; the head was allowed to move freely and was not in a fixed position. Mice were anesthetized with a combination of ketamine (30 mg/kg, intraperitoneal [IP]) and xylazine (5 mg/kg, IP) before each blast or sham blast, and were placed on a heating pad immediately following blast injury to prevent hypothermia and facilitate recovery from general anesthesia. Xylazine anesthesia was reversed with Yohimbine chloride (0.001 mg/g, IP) to aid in recovery from anesthesia. Control mice underwent an identical process in all respects except that they did not receive a blast exposure when placed in the chamber. Both blasted and sham mice were given an intraperitoneal injection (0.1mL/20g body weight) of buprenorphine (0.003 mg/mL) immediately after the blast or sham-blast, respectively.

**Figure 1.**
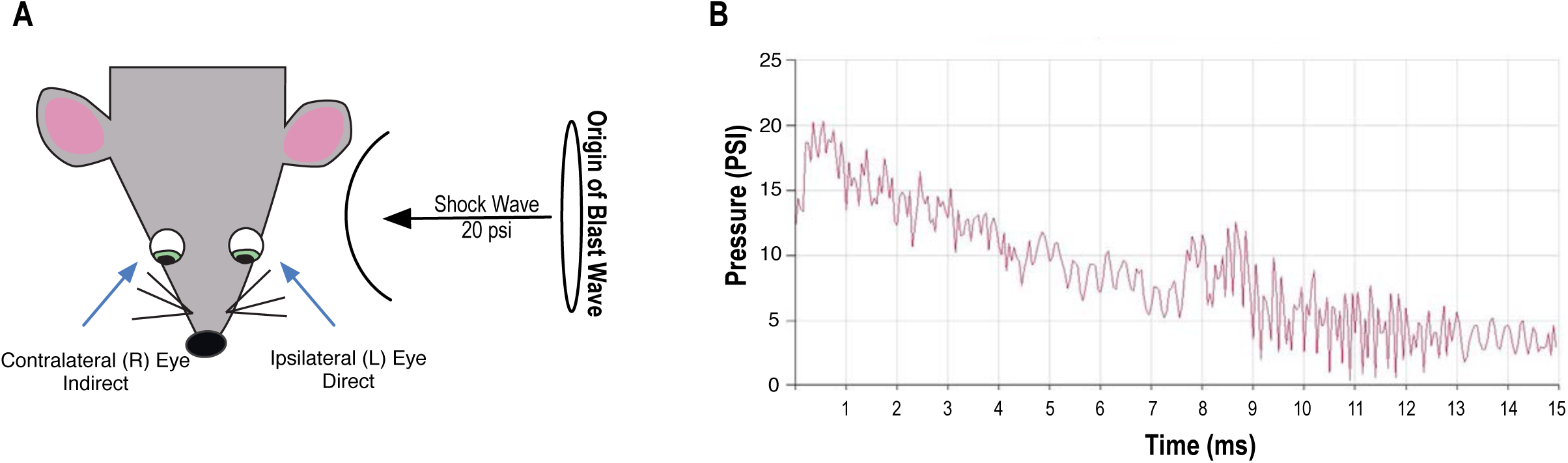
Schematic representation of blast exposure. Exposure to blast wave pressure was conducted as previously described (Mohan, Kecova et al. 2013). Animals were placed lateral to the shock tube axis, 30 cm from the origin of the blast wave (Mylar membrane), with the left (L) side of the head (ipsilateral eye) facing the blast wave. Control mice were restrained the same way, but were not exposed to the blast wave (A). A representative tracing of a 20 ± 0.2 psi (137.8 ± 1.3 kPa, mean ± SEM) blast wave (B). R, Right.

### RNA isolation, cDNA Preparation, and Real Time Quantitative Polymerase Chain Reaction (RT-qPCR)

Total RNA was extracted from sham or blast brain regions using TRIzol (Invitrogen, Carlsbad, CA) as per the manufacturer’s instructions. Retinas were lysed with a homogenizing pestle and filtered through QiaShredder columns (Qiagen, Chatsworth, MD, USA). RNA was then extracted with an RNeasy Mini Kit (Qiagen) according to the manufacturers instructions. For cDNA preparation of brain regions and retinas, respectively, 2000 ng or 1000 ng of total RNA was reversed transcribed using random hexamers and SuperScript III Reverse Transcriptase (Invitrogen). Amplified cDNAs were diluted 1:10 and 1:15 for retina and brain samples, respectively, in ultrapure water and subjected to real-time PCR using an Applied Biosystems Model 7900HT with TaqMan Universal PCR Mastermix (Applied Biosystems) with the following probes: *IL-1β* (Mm00434228_m1), *IL-1α* (Mm00439620_m1), *IL-6* (Mm00446190_m1), *TNF-α* (Mm00443258_m1), and *GAPDH* (4308313). Biologic samples were run in triplicate. Relative mRNA levels of target genes were normalized to the endogenous control, *GAPDH*, using the comparative cycle threshold method. Results were expressed as fold difference from sham controls. Column statistics were run to determine if values were normally distributed. For those with normal distribution, a Student’s *t*-test was used, otherwise a Mann-Whitney U Test was utilized.

### Immunohistochemistry (IHC)

Blast-exposed and sham mice were deeply anesthetized with 200 mg/kg ketamine and 20 mg/kg xylazine and perfused transcardially with 0.01 mol/L phosphate-buffered saline (PBS) followed by 4% paraformaldehyde in 0.01 mol/L PBS. Ipsilateral globes were post-fixed in 4% paraformaldehyde for 1 hour at room temperature.

For cross-section, the lens was removed and globes were subjected to a sucrose gradient (10%, 20%, and 30% in 0.01 mol/L PBS), embedded in OCT, and frozen using dry ice. Sagittal sections (8 μm thick) of the ipsilateral retina were collected onto superfrost plus glass slides. OCT-embedded sections were hydrated for 15 minutes with tris-buffered saline with 0.05% Tween 20, followed by 45 minutes of permeabilization with 0.3% triton in PBS. Sections were blocked for 1 hr with 3% goat serum/0.1% Triton X-100 (cat. no. T8532; Sigma-Aldrich, St. Louis, MO) in PBS. Primary antibodies against glial fibrillary acidic protein (GFAP; 1:400; cat. no. Mab360; Millipore, Burlington, MA) and IBA-1 (1:500; cat. no. 019-1941; Wako Chemicals USA, Inc., Richmond, VA) were diluted in blocking solution and costained with fluoresceinated Griffonia simplicifolia type 1 Isolectin B4 (Alexa Fluor 594 conjugated, 1:100, cat. no. 121413; Life Technologies, Grand Island, NY, USA) in 1 mM CaCl2 in PBS overnight at 4°C. Tissues were washed with PBS (3 × 5 minutes), and incubated with corresponding secondary antibodies GAR488 (1:300; cat. no. A11070; Thermo Fisher Scientific, Waltham, MA) and GAM647 (1:300; cat. no. A-32728; Thermo Fisher Scientific) with Isolectin B4 (1:100) in 3% goat serum/0.1% triton in PBS for 1 hour at room temperature. Samples were washed with PBS (3 × 5 min), incubated with DAPI (4ug/ml; cat. no. D9564; Sigma-Aldrich) for 10 min at room temperature, and washed with PBS (3 × 5 min).

For whole-mount IHC, retinal dissection was followed by permeabilization with 0.3% Triton X-100 in PBS for 1 hour at RT and antigen retrieval treatment (10 mM sodium citrate, pH=6 with 0.05% Tween 20) at 90°C for 10 minutes. Retinas were blocked overnight with 3% goat serum/0.1% triton in PBS at 4°C, and then incubated for 1.5 hours with Background Buster (cat. no. NB306; Innovex Biosciences Inc., Richmond, CA). Retinas were incubated with primary antibody as described above overnight at 4°C, washed with PBS (3 × 5 min), and incubated with corresponding secondary antibodies for 1 hour at room temperature. All slides were mounted with ProLong Gold Antifade Mountant (cat. no. P36934; Thermo Fisher Scientific) followed by imaging with confocal microscopy (SP8 confocal, Leica Microsystems, Wetzlar, Germany).

### Quantification of IBA-1 Immunohistochemistry Staining

Flat-mounted retinas were stained as described above. Two nonoverlapping images were collected for each of 4 retinal petals, yielding a total of 8 images per retina, and were taken midway between the outer edge of the petal and the optic nerve head (**Fig. 5A**). Images were taken using a X25 water-immersion objective with consistent microscope settings to eliminate variation from one sample to the next. Quantitative analysis of area and cell counts were performed using Image J Imaging Software (National Institutes of Health, Bethesda, MD) (Rueden, Schindelin et al. 2017) by an observer masked to the origin of the sections. Values from the 8 images were averaged, producing one value per retina. Results were expressed as mean ± SEM and a Student’s *t*-test was performed. A value of *P*< 0.05 was considered significant.

### Treatment with Anakinra

Blast and sham mice were randomly assigned to recieve intraperitoneal injections once daily with a dose of 100 mg/kg or an equal volume of sterile 0.9% saline. Anakinra (67mg/mL; Sobi) was diluted in sterile 0.9% saline. Treatment was initiated one week before blast injury and continued for 3 weeks post-injury. Users were blinded to the treatments for the entirety of the study.

### Pattern-Evoked Electroretinography

Pattern-evoked electroretinography (PERG) was used to objectively measure the function of retinal ganglion cells (RGCs) by recording the amplitude of the PERG waveform following TBI. Mice were anesthetized with a combination of ketamine (0.05-0.06 mg/g, IP), xylazine (0.008-.01 mg/g, IP), and acepromazine (0.004-.005 mg/g, IP), and then placed on a heated recording table to maintain body temperature. Neutral position PERG responses were evoked using alternating, reversing, and black and white vertical stimuli delivered on a monitor with a Roland Consult ERG system (Roland Consult, Brandenburg, Germany). To record the PERG response, commercially available stainless steel subdermal electrodes (Ambu, Ballerup, Denmark) were placed in the snout. A reference needle electrode was placed medially at the base of the head, and a ground electrode was placed at the base of the tail to complete the circuit. Each animal was placed at the same fixed position in front of the monitor to prevent recording variability due to animal placement. Stimuli (18° radius visual angle subtended on full field pattern, 2 reversals per second, 372 averaged signals with cut off filter frequencies of 1 to 30 Hz, 98% contrast, 80 cd/m^2^ average monitor illumination intensity (Jorvec, Miami, FL, USA)) were delivered under mesopic conditions without dark adaptation to exclude the possible effect of direct photoreceptor-derived evoked responses. A diffuser placed over the pattern on the monitor also did not elicit a measurable evoked potential, further ensuring that the electrical responses were elicited from RGCs. The PERG response was evaluated by measuring the amplitude (peak to trough) of the waveform, as we have described previously (Mohan, Harper et al. 2012, Mohan, Kecova et al. 2013). The PERG response was recorded at baseline, and 4 weeks following blast injury. Statistical significance was determined by one-way analysis of variance (ANOVA) followed by Dunnet’s post test for multiple comparisons using the sham-saline group as the baseline.

### Spectral-Domain Optical Coherence Tomography (SD-OCT)

SD-OCT analysis was performed at baseline and 4.5 weeks post injury using a Spectralis SD-OCT (Heidelberg Engineering, Vista, CA) imaging system and a 25 dipoter (D) lense for mouse ocular imaging (Heidelberg Engineering). Mice were anesthetized with a combination of ketamine (0.03 mg/g, IP) and xylazine (0.005 mg/g, IP) and placed on a heating pad to maintain body temperature. Pupils were dilated using a 1% tropicamide solution and the cornea was moisturized with saline. Volume scans (49 line dense array) positioned directly over the optic nerve head were performed to quantify the RGC complex thickness, which includes RGC bodies, axons, and dendrites. Scans approximately 100 um from the edge of the optic nerve head were analyzed by a masked observer, excluding blood vessels from the RGC complex thickness calculation. Statistical significance was determined by one-way analysis of variance (ANOVA) with Dunnet’s post test for multiple comparisons.

### Histology and Microscopy of Optic Nerves

Blast-exposed and sham-blast mice were deeply anesthetized with carbon dioxide, lightly perfused with normal saline followed by 4% paraformaldehyde and euthanized by decapitation. Optic nerves were collected and processed as previously described (Mao, Hedberg-Buenz et al. 2011). In brief, optic nerves were dissected from heads and drop fixed in in half-strength Karnovsky’s fixative (2% paraformaldehyde, 2.5% gluteraldehyde in 0.1 M sodium cacodylate) at 4°C for 16 hours. Nerves were rinsed in 0.1 M Na cacodylate buffer, post-fixed with 1% osmium tetroxide, dehydrated in graded acetone (30%-100%), infiltrated in graded resin (33%, 66%, and 100%; Eponate-12; Ted Pella, Redding, CA) diluted in propylene oxide, embedded in fresh 100% resin, and then polymerized in a 65°C oven. Semithin (1-µm) cross sections were cut, transferred to glass slides, stained with 1% paraphenylenediamine (PPD), and mounted to glass slides (Permount, Fisher Scientific, Pittsburgh, PA). Light micrograph images were obtained with an Olympus BX-52 microscope (Cedar Valley, PA) at total magnifications of 100X and 1000X.

### Grading of Optic Nerve Damage

Assessing levels of damage in optic nerves was performed using a 3-level grading scale (1-none to mild damage, 2-moderate, or 3-severe) as previously described (Anderson, Libby et al. 2005, Libby, Anderson et al. 2005). Microscopy slides containing mounted optic nerve cross sections stained with PPD were assessed by two independent investigators that were blinded to both the identity of the nerves and the scores given by the other investigator. Investigators assigned the same score for 91% of the nerves. In the cases of damage grade disagreement, a third investigator, blinded to both the identity of nerves and scores given by the first two investigators, assigned a third damage grade and the most common grade (amongst all three investigators) was used in the final grading for each nerve specimen. Representative light micrographs were obtained from slides using a light microscope (BX-52, Olympus) at total magnifications of 100X and 1000X.

### Experimental Design and Statistical Analysis

Column statistics were run to determine if values were normally distributed for all quantitative experiments. For the RT-qPCR data and quantification of immunohistochemistry at 4 hours, 24 hours, and 1 week, a Student’s *t*-test was used if data were normally distributed, otherwise a Mann-Whitney U Test was utilized. Statistical significance for the PERG and OCT analysis was determined by one-way analysis of variance (ANOVA) followed by Dunnet’s post test for multiple comparisons using the sham-saline group as the baseline. A value of *P*< 0.05 was considered significant. All the statistical data were presented as mean ± SEM and were done using GraphPad Prism 7 for Macintosh (La Jolla, CA, USA).

## Results

### Repeated bTBI Causes Increased Expression of Inflammatory Cytokines in the Retina and Brain

As robust neuroinflammation occurs in the brain following several types of TBI (Simon, McGeachy et al. 2017), we first examined the mRNA levels of classic pro-inflammatory cytokines in the retina and brain after bTBI. At 4 hours post repeated bTBI, there was an acute increase in retinal inflammatory cytokine mRNA when normalized to sham values, with increased expression of *IL-1β* (*P*<0.0001), *IL-1α* (*P*=0.003), *TNFα* (*P*=0.0001), and *IL-6* (*P*<0.0001) in the ipsilateral retinas of blast-injured mice (**Fig 2**). Increases in *IL-1β, IL-1α, TNFα*, and *IL-6* were seen in brain tissue of mice exposed to blast injury in a region-dependent manner 4 hours after injury. *IL-1β* was increased in all regions analyzed: cerebellum (CB), hippocampus (HIPP), extra forebrain (XFB, tissue block contains striatum, thalamus, hypothalamus, and visual white matter tracts), and brain stem (BS) samples of bTBI mice (*P*<0.0001, *P*=0.0085, *P*=0.0046, *P*=0.0002, respectively; **Fig 3A**). *IL-1α* was increased in CB and BS (*P*=0.0046, *P*=0.0085, respectively; **Fig 3B**), *TNFα* was increased in XFB and BS (*P*=0.0103, *P*<0.0001, respectively; **Fig 3C**), and *IL-6* was increased in CB (*P*=0.0489; **Fig 3D**) of blast-injured samples. While inflammatory mRNA expression varied between regions and also between individual samples, these increases are indicative of acute inflammatory changes overall in the brain in addition to those seen in the retina.

**Figure 2.**
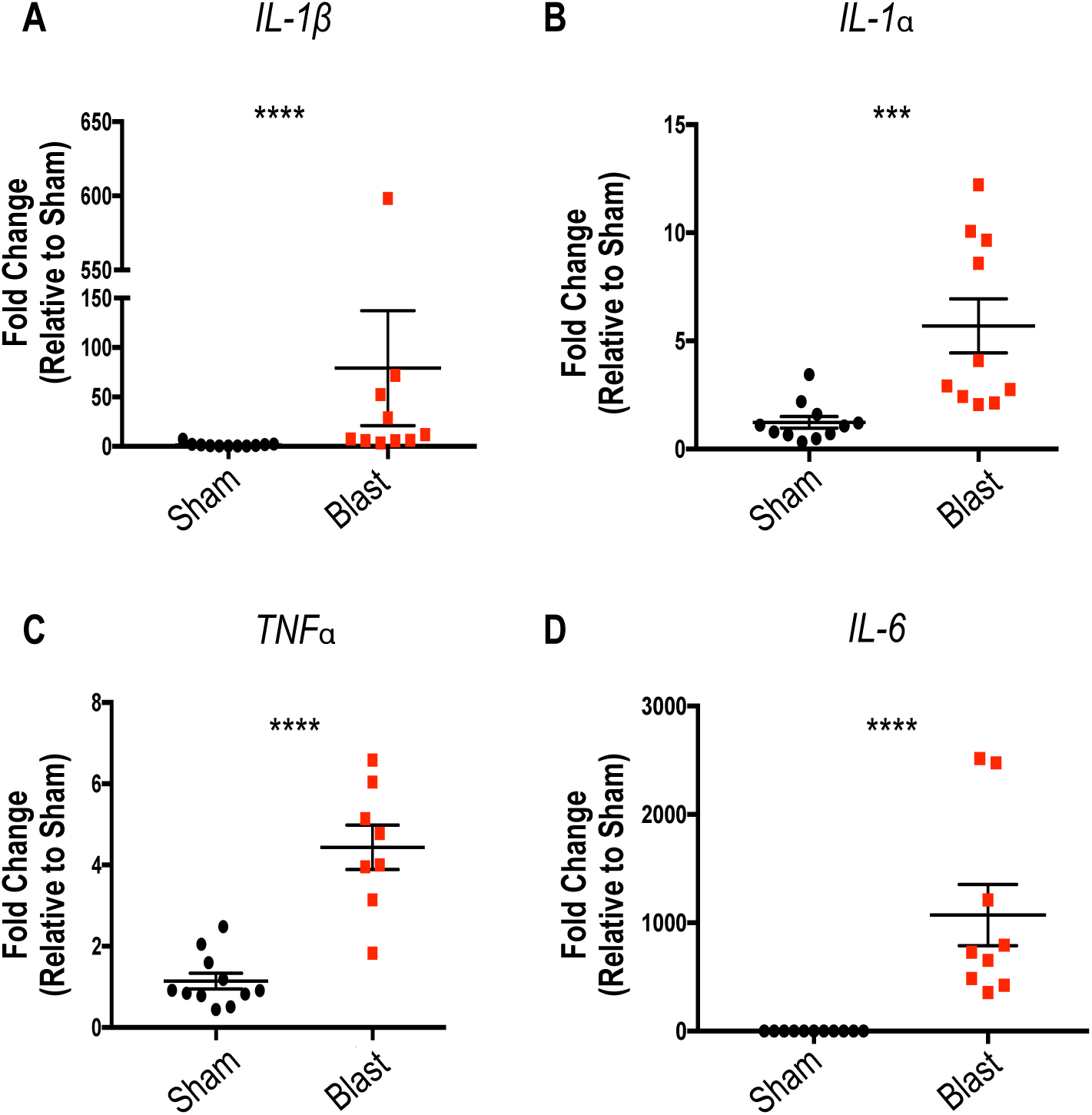
Retinal expression of inflammatory markers is increased 4 hours post repeated bTBI. Quantitative PCR measuring mRNA levels relative to *GAPDH*. Increased expression of *IL-1β* (**A)**, *IL-1α* (**B)**, *TNFα* (**C**), and *IL-6* (**D**) in the ipsilateral retinas of blast-injured mice was seen when normalized to sham values. Student’s *t*-Test or Mann-Whitney U test based on distribution of data. *** *P*<0.001; **** *P*<0.0001.

**Figure 3.**
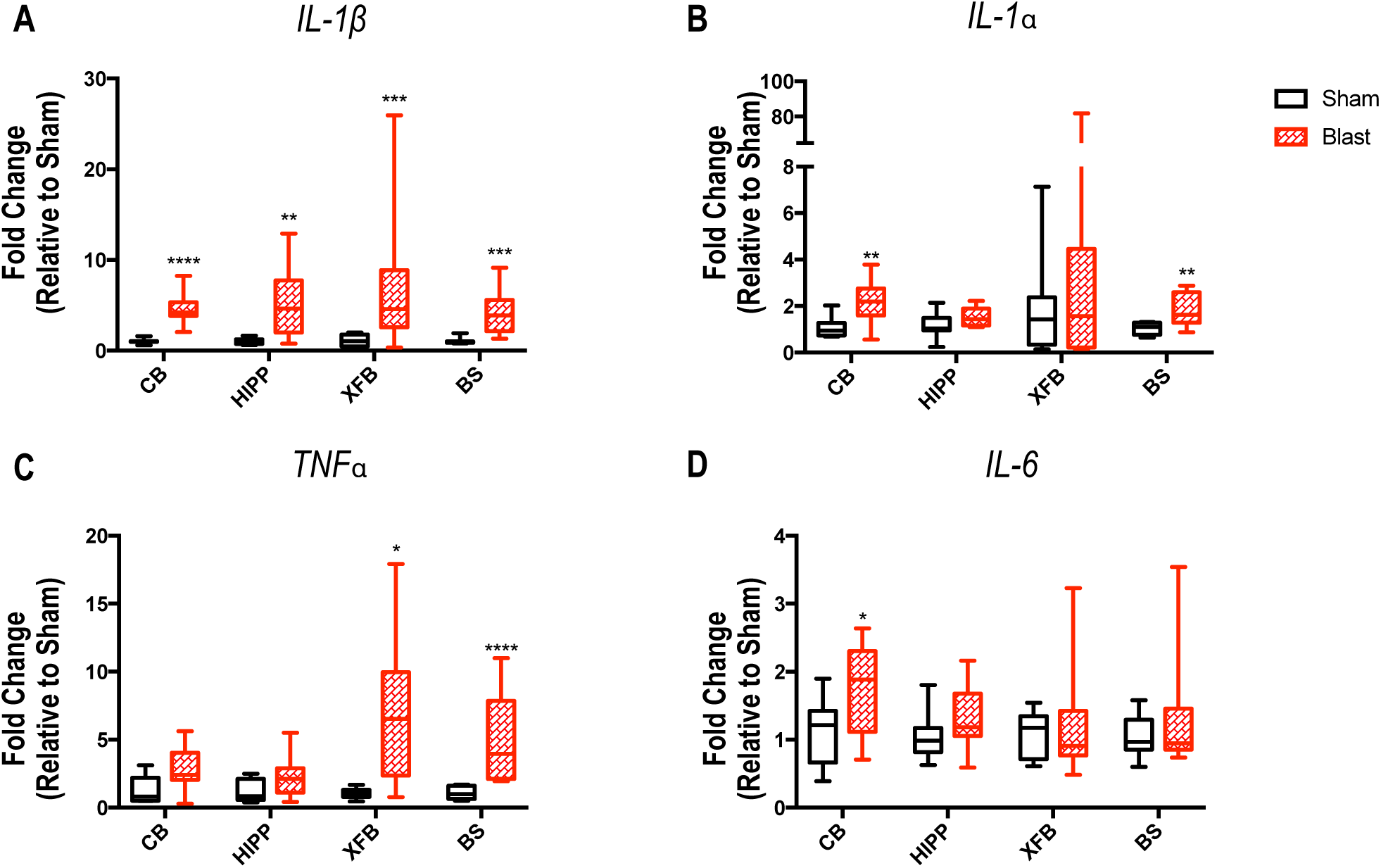
Inflammatory cytokine expression in brain tissue demonstrates regional changes 4 hours following repeated bTBI. Expression of *IL-1β* (**A**), *IL-1α* (**B**), *TNFβ* (**C**), and *IL-6* (**D**) was evaluated by quantitative PCR in ipsilateral cerebellum (CB), hippocampus (HIPP), extra forebrain (XFB), and brainstem (BS). mRNA levels relative to the housekeeping gene *GAPDH*. Data are expressed as fold change in gene expression relative to sham and are presented as box and whiskers plots; the box extends from 25th to 75th percentiles, the line represents the median, and the whiskers extend from smallest to largest value. Student’s *T*-test or Mann-Whitney U test based on distribution of data. \**P*<0.05; \*\**P*<0.01; \*\*\**P*<0.001; \*\*\*\**P*<0.0001.

### Repeated bTBI Induces Cellular Activation in the Retina due to Injury

The mammalian retina contains three types of glial cells; microglia are the resident phagocytes of the retina along with two forms of macroglial cells, astrocytes and Müller (radial glial) cells, which carry out neuronal support functions in the retina (Bringmann, Pannicke et al. 2006, Cuenca, Fernandez-Sanchez et al. 2014). TBI triggers rapid activation of resident microglia and macroglia, all of which can be activated by and continue to produce inflammatory cytokines (Allan, Tyrrell et al. 2005). The prevalence and morphology of retinal microglia in the retinal nerve fiber layer (RNFL) and retinal ganglion cell (RGC) layer in response to blast injury was assessed 4 hours, 24 hours, and 1 week after injury using the marker IBA-1. In sham mice, IBA-1 immunoreactivity showed ramified microglia with thin irregular processes, reflecting their resting state (**Fig 4A-C**). Ispilateral blast-injured retinas demonstrated hyper-ramified and bushy microglia with thicker processes and swollen cell bodies at all three timepoints (**Fig 4D-F**), characteristic morphologic changes indicative of activation due to injury (Crews and Vetreno 2016).

**Figure 4.**
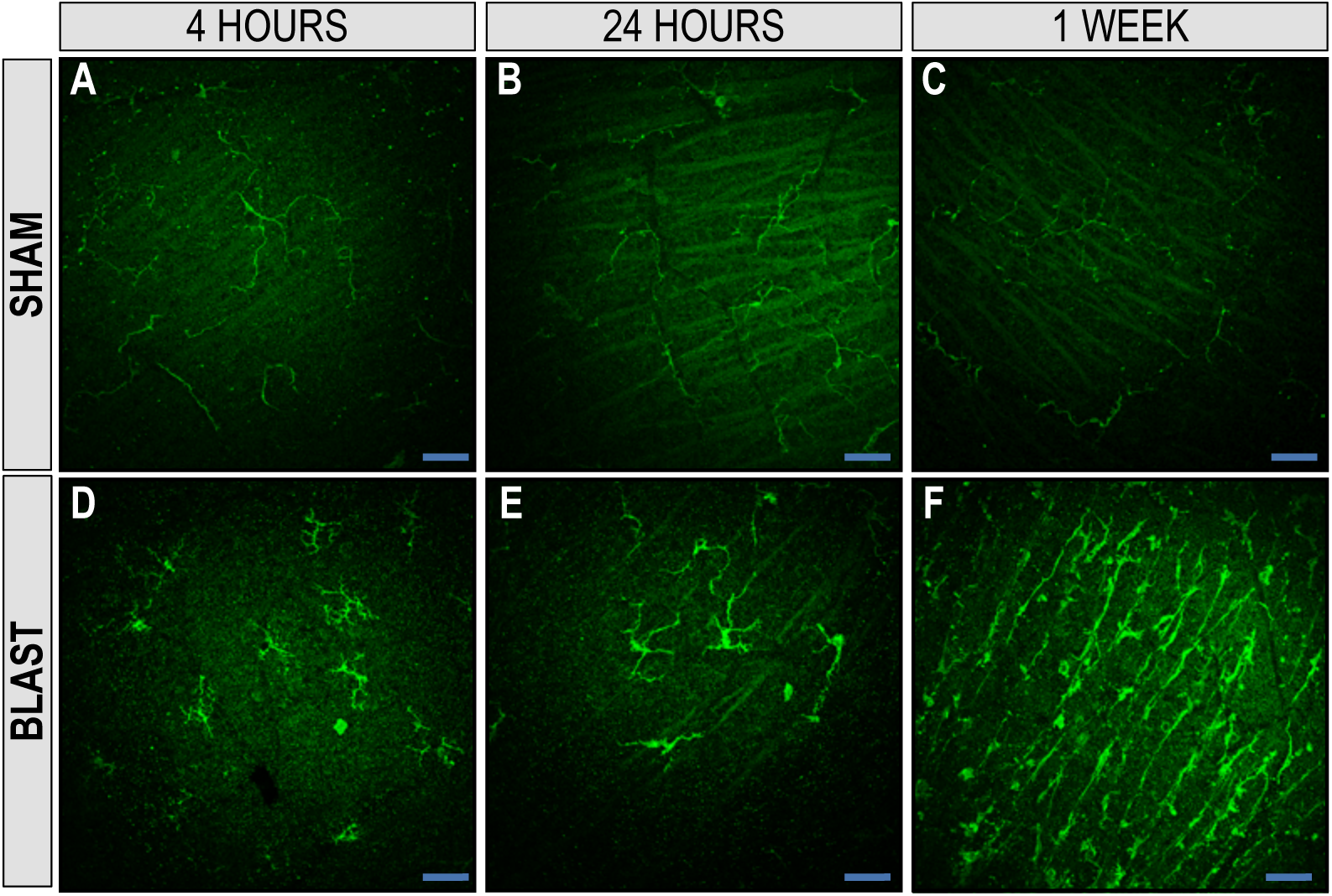
Microglia are activated in retinas of blast-injured animals. IBA-1+ microglia in the RNFL and RGC layer of retinal whole-mounts exposed to blast injury. Retinas from sham animals had a ramified morphology at 4 hr, 24 hr, and 1 week (**A, B**, and **C**, respectively). At 4 hr, 24 hr, and 1 week post injury (**D, E**, and **F**, respectively), blast-injured retinas demonstrated hyper-ramified and bushy microglia, suggesting activation due to injury. Original magnification, X25; en face view with the RNFL facing up. Scale bar: 50 µm. (RNFL: retinal nerve-fiber layer; RGC: retinal ganglion-cell layer).

For the 24 hours and 1 week time points, IBA-1 quantification (average of 8 sample regions/retina; **Fig 5A**) was carried out to evaluate the percent area of fluorescence of IBA-1 possitive microglia and the number of cell bodies present in the RNFL and RGC layer in sham and blast mice (representative images **Fig 5B, C**, respectively). At 24 hours the percent area of fluorescence (**Fig 5D**, bTBI=3.15 ± 0.3044; sham=1.93 ± 0.28; *P*=0.0163) and the number of cell bodies (**Fig 5G**, bTBI=7.707 ± 0.5575; sham=4.371 ± 0.6637; *P*=0.0046) was significantly increased in the retinas of blast mice (**Fig 5F, I**) when compared to sham (**Fig 5E, H**; arrow heads indicate cell bodies), respectively. This same finding was seen 1 week post injury, with the percent area of fluorescence (**Fig 5J**, bTBI=5.065 ± 1.318; sham=1.869 ± 0.04373; *P*=0.0043) and the number of cell bodies (**Fig 5M**, bTBI=12.97 ± 2.797; sham=5.019 ± 0.281; *P*=0.0122) significantly increased in the retinas of blast mice (**Fig 5L, O**) when compared to sham (**Fig 5K, N**), respectively.

**Figure 5.**
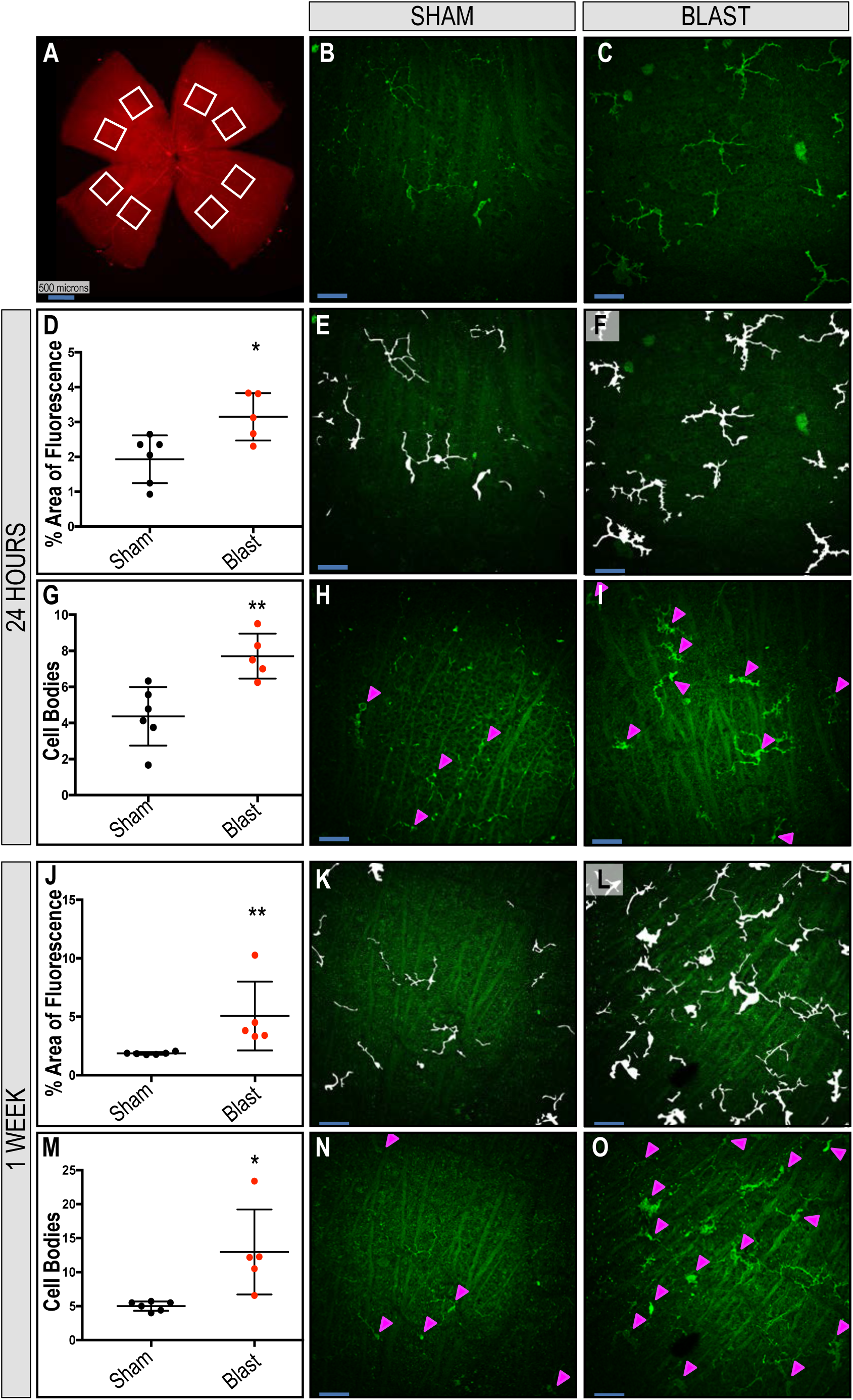
Increased immunoreactivity and distribution of IBA-1 positive microglia is found in retinas exposed to blast. Eight regions sampled per ipsilateral retina (**A**). Quantification shows an increase in total fluorescence area (**D, J**) and cell body counts (**G, M**) after blast in the RNFL and RGC layer of whole-mounted retinas at 24 hr and 1 wk, respectively. Representative images (**B, C**) and mask overlays of area quantified (**E, H, K, M** and **F, I, L, O**) for sham and blast-injured mice, respectively. Arrow heads indicate cell bodies (**F, I**). Student’s *t*-Test or Mann-Whitney U test based on distribution of data. \**P*<0.05; \*\**P*<0.01. Data expressed as means ± SEM. n=5-6 mice per group. Original magnification, X25; en face view with the RNFL facing up. Scale bar: 50 µm unless otherwise noted. (RNFL: retinal nerve-fiber layer; RGC: retinal ganglion-cell layer).

Astrocytes and Müller glia, the macroglia cells of the retina, carry out many homeostatic functions in order to maintain the retinal extracellular environment and health. Müller glia have processes that project radially throughout the entire span of the retina, aiding in the early detection of retinal stress, and provide limits at the outer and inner limiting membrane. In the healthy retina, GFAP is expressed abundantly in astrocytes in the RNFL and is localized to the end feet of Müller glia at the retina’s inner limiting membrane (Bringmann, Pannicke et al. 2006). However, in response to retinal injury or stress, GFAP immunoreactivity is increased in Müller glial processes and can extend throughout the retina (Lewis and Fisher 2003). We assessed the distribution of GFAP immunoreactivity in retinal cross sections to detect activation of Müller glial cells after blast exposure. 24 hours after injury, GFAP immunoreactivity was limited to astrocytes in the RNFL in ipsilateral retinas of both sham and blast-injured mice (**Fig 6A, B**, respectively). One week after blast exposure, the distribution of GFAP immunoreactivity remains in the RNFL of sham mice (**Fig 6C**), however, it is consistently increased in bTBI retinas, spanning the retina radially from the RNFL to the outer nuclear layer (ONL; **Fig 6D**). A 3D reconstruction of a z-stack taken of whole-mounted retinas were rotated to demonstrate that GFAP immunoreactivity is limited to the RNFL of sham mice (**Fig 6E**), while spanning the retina in blast mice (**Fig 6F**) as a response to injury.

**Figure 6.**
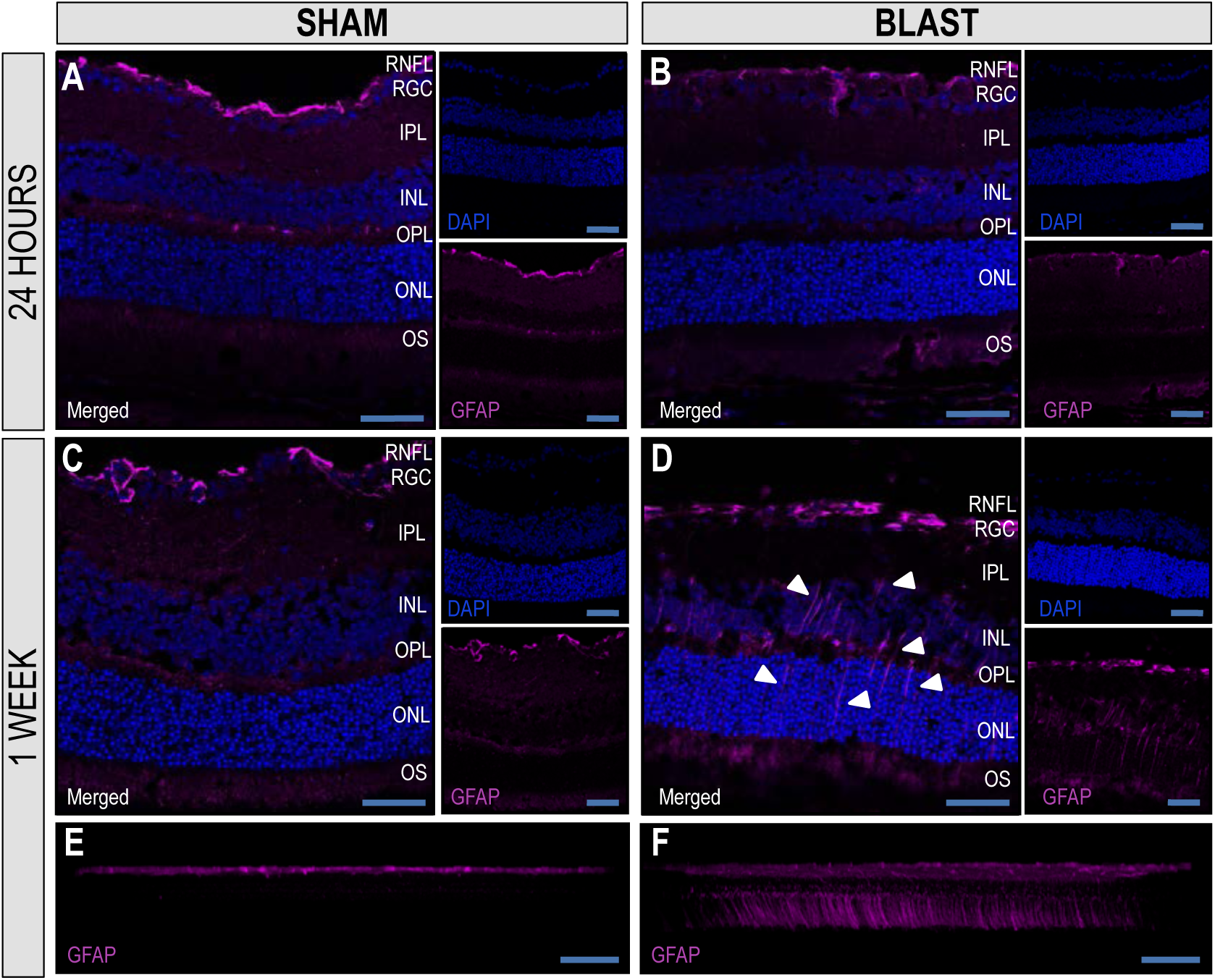
Extended distribution of GFAP in Müller glia in retinas ipsilateral to blast exposure suggests increased damage. GFAP localizes to the retinal nerve fiber layer (RNFL) of retinas of sham and blast mice 24 hours after injury (**A, B, respectively**). One week after blast exposure, the distribution of GFAP immunoreactivity remains in the RNFL of sham mice (**C**), but increases in Müller glia spanning the retina from the RNFL to the outer nuclear layer (ONL) in blast mice (**D**). 3D reconstruction of a z-stack of 0.5 μm confocal sections at X25 magnification rotated to demonstrate GFAP reactivity limited to the RNFL of sham mice (**E**) and spanning throughout the retina in blast mice (**F**). (RGC: retinal ganglion-cell layer; IPL: inner plexiform layer; INL: inner nuclear layer; OPL: outer plexiform layer; ONL: outer nuclear layer: OS: photoreceptor outer segments).

**Figure 7.**
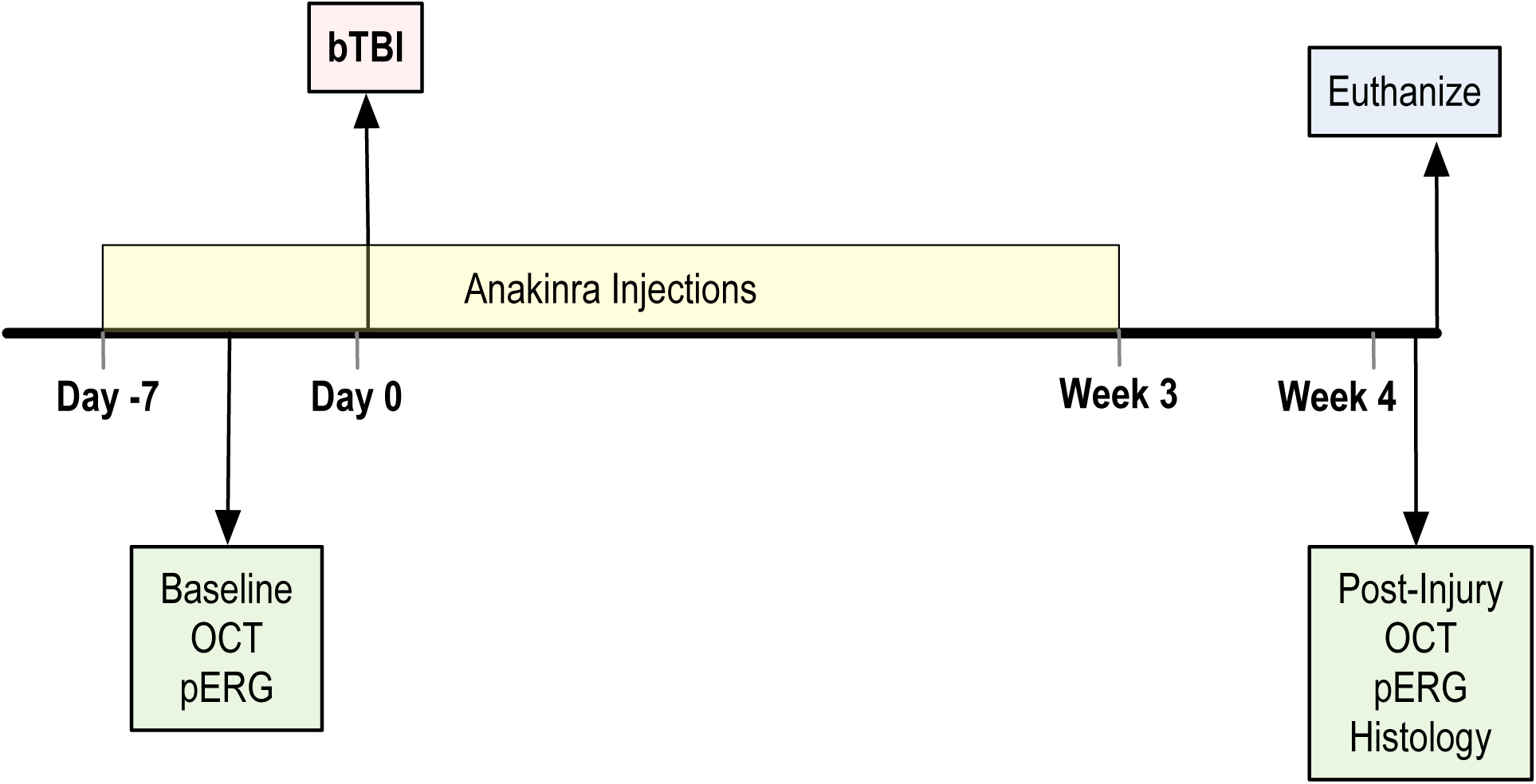
Experimental design of anakinra treatment. Mice were given injections of either saline or anakinra one week before injury and three weeks post. Optical coherence tomography (OCT) and pattern electroretinogram (PERG) were conducted before injury as a baseline and four weeks post, followed by euthanasia and tissue collection for histological analysis of retinal and optic nerve tissue.

### Anakinra Reduces Cellular Activation after bTBI in the Retina

To determine if inflammatory blockade could decrease retinal damage after bTBI, we treated both sham and blast mice with either saline or anakinra (**Fig. 7**). We carried out quantification of the percent area of fluorescence (**Fig 8A-D**) and number of cell bodies (**Fig 8F-I**) of IBA-1 positive retinal microglia in the RNFL and RGC layer in response to blast injury 4 weeks after injury for our four conditions: sham-saline (SS), blast-saline (BS), sham-anakinra (SA), and blast-akaninra (BA), respectively. When compared to the sham-saline group as baseline, the blast-saline group was the only condition that had significantly higher percent area of fluorescence (**Fig 8E**, SS=1.165 ± 0.1758; BS=2.349 ± 0.3969; SA=1.022 ± 0.1853; BA=1.912 ± 0.1937; *P*=0.0063) and a significantly increased number of cell bodies (**Fig 8J**, SS=6.234 ± 0.5487; BS=13.03 ± 1.669; SA=5.063 ± 0.727; BA=8.717 ± 1.385; *P*=0.0007), suggesting that anakinra treatment prevented microglia activation after blast injury.

**Figure 8.**
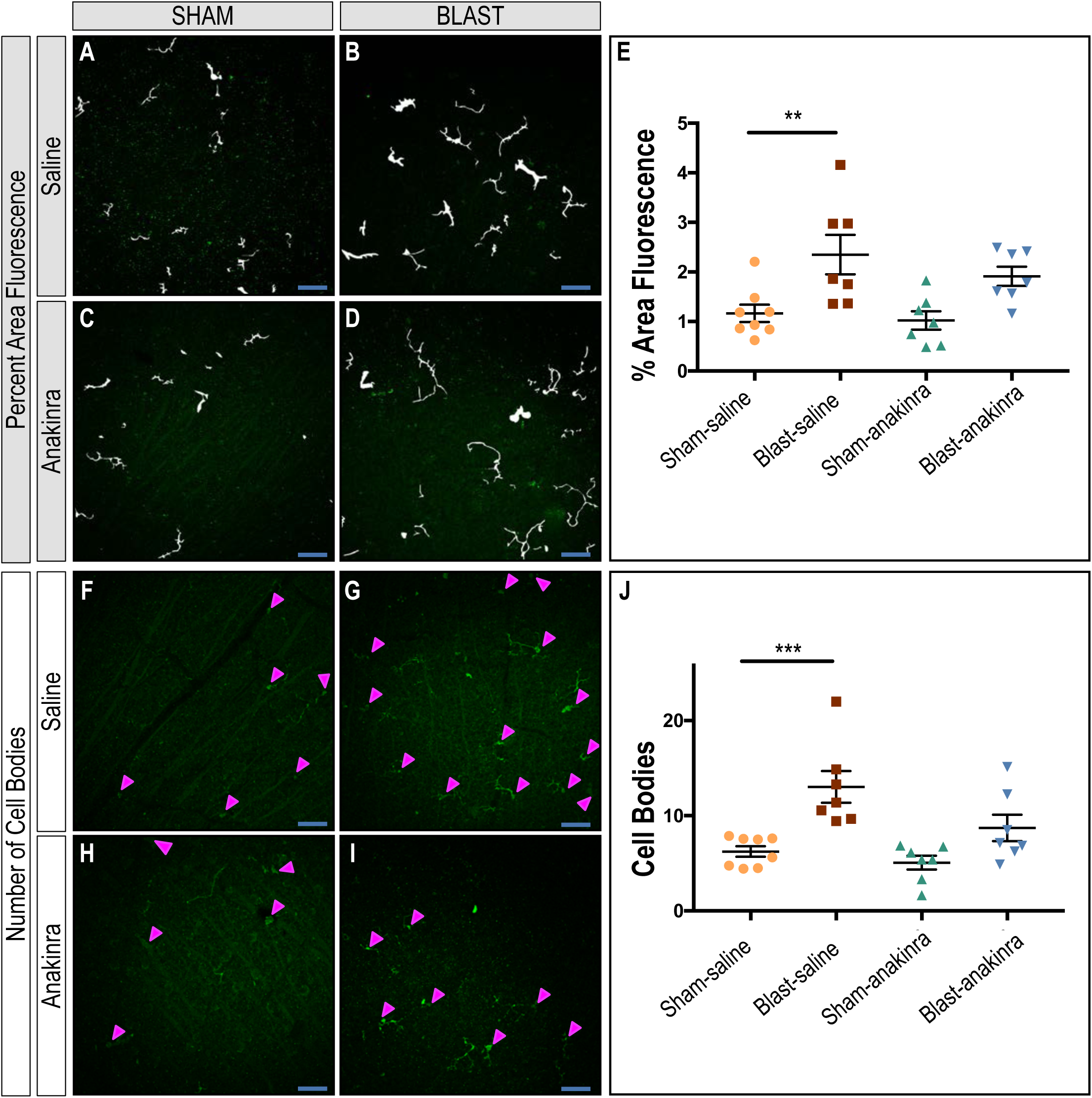
Anakinra treatment reduces microglial activation in the retina after blast. Mask overlays of the fluorescent area quantified (**A, B, C, D**) and arrow heads indicating cell bodies (**F, G, H, I**) for ipsilateral retinas of sham-saline, blast-saline, sham-anakinra, and blast-anakinra groups, respectively. Quantification shows an increase in total fluorescent area **(F)** and cell body counts **(J)** in the RNFL and RGC layer of whole-mounted retinas from blast-saline mice. No significant difference was found comparing the sham-anakinra and the blast-anakinra groups to the sham-saline. Significant difference compared to sham-saline group using one-way ANOVA with Dunnett’s posttest (\*\**P*<0.01, \*\*\**P*<0.001). Data expressed as means ± SEM. Eight regions sampled per retina; n=7-8 mice per group. Original magnification, X25; scale bar: 50 µm. (RNFL: nerve-fiber layer; RGC: retinal ganglion-cell layer).

We also assessed the effects of anakinra on Müller glia activation via the distribution of GFAP immunoreactivity in retinal cross sections 4 weeks after blast exposure in three separate mice per condition. Both the sham-saline (**Fig 9A-C**) and sham-anakinra (**Fig 9D-F**) groups demonstrated GFAP localization limited to the RNFL. Retinas from the blast-saline mice exhibited increased GFAP immunoreactivity extending from the RNFL into deeper retinal layers, indicative of glial activation due to retinal stress (**Fig 9G-I**). However, the GFAP immunoreactivity was consistently visibly decreased in retinas from blast-anakinra mice (**Fig 9J-L**), suggesting that anakinra at least partially prevented Müller glia activation in blast-injured retinas.

**Figure 9.**
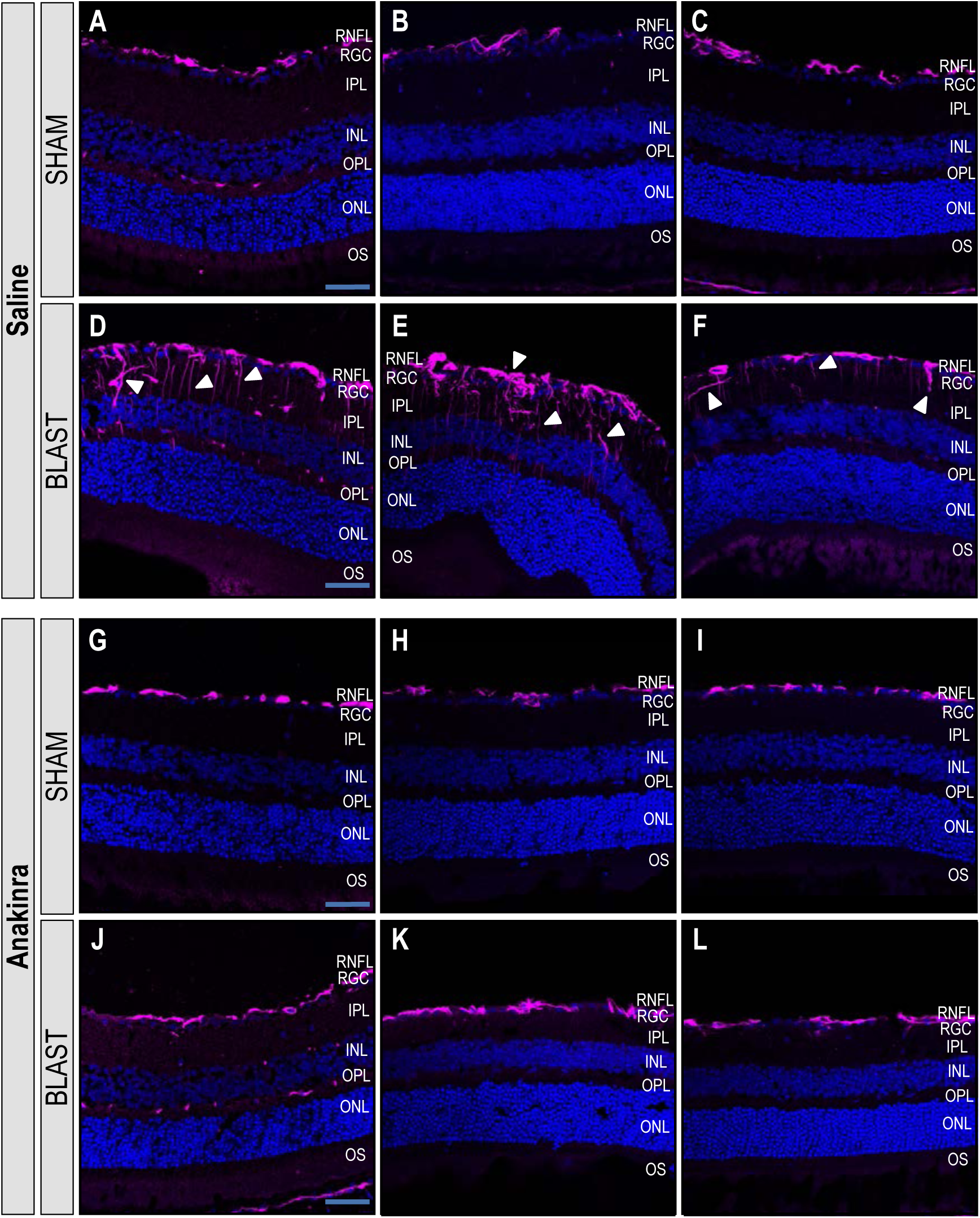
Anakinra treatment prevents Müller glia activation in the retina after blast. Distribution of GFAP in retinas ipsilateral to blast exposure 4 weeks after injury. Images of areas with the highest GFAP reactivity shown in retinal cross sections of three individual mice per group. GFAP localizes to the retinal nerve fiber layer (RNFL) of retinas of sham-saline (**A-C**) and sham-anakinra (**D-F**) mice 4 weeks after blast injury. Retinas from blast-saline mice demonstrate increased GFAP immunoreactivity in Müller glia spanning the retina from the RNFL into deeper layers (**G-I**). However, blast-anakinra mice show decreased GFAP immunoreactivity overall (**J-L**) when compared to the injury group only given saline. (RGC: retinal ganglion-cell layer; IPL: inner plexiform layer; INL: inner nuclear layer; OPL: outer plexiform layer; ONL: outer nuclear layer: OS: photoreceptor outer segments).

### Anakinra Prevents Blast-Induced RGC Damage after bTBI

While anakinra decreased retinal glial activation after bTBI, we wanted to assess the changes in RGC structural and functional outcomes after blast with this treatment. RGC cell death can result in a loss of their axons compromising the optic nerve and resulting in visual field loss. We conducted histological analysis of optic nerves stained with paraphenylenediamine (PPD) to highlight myelinated axons. The level of neurodegeneration was based on a grading system, with Grade 1 as healthy-mildly damaged (**Fig 10A, B**), Grade 2 as moderate damage (**Fig 10C, D**), and Grade 3 as severe damage (**Fig 10E, F**) when evaluating PPD-positive and infilled damaged axons (arrow heads) and glial scar formation (asterisks) adjacent to glial cell nuclei. The sham-saline and sham-anakinra optic nerves were most frequently healthy with Grade 1 damage (SS= Grade 1: 90.9%, Grade 2: 9.1%; SA= Grade 1: 90%, Grade 2: 10%), while the blast-saline group was more frequently graded as more damaged (Grade 2: 27.27%, Grade 3: 72.72%). However, treatment with anakinra resulted in shifting the grading severity to lesser damage levels (Grade 1: 25%, Grade 2: 25%, Grade 3: 50%), suggesting that anakinra prevents some of the damage to the optic nerve that occurs due to bTBI.

**Figure 10.**
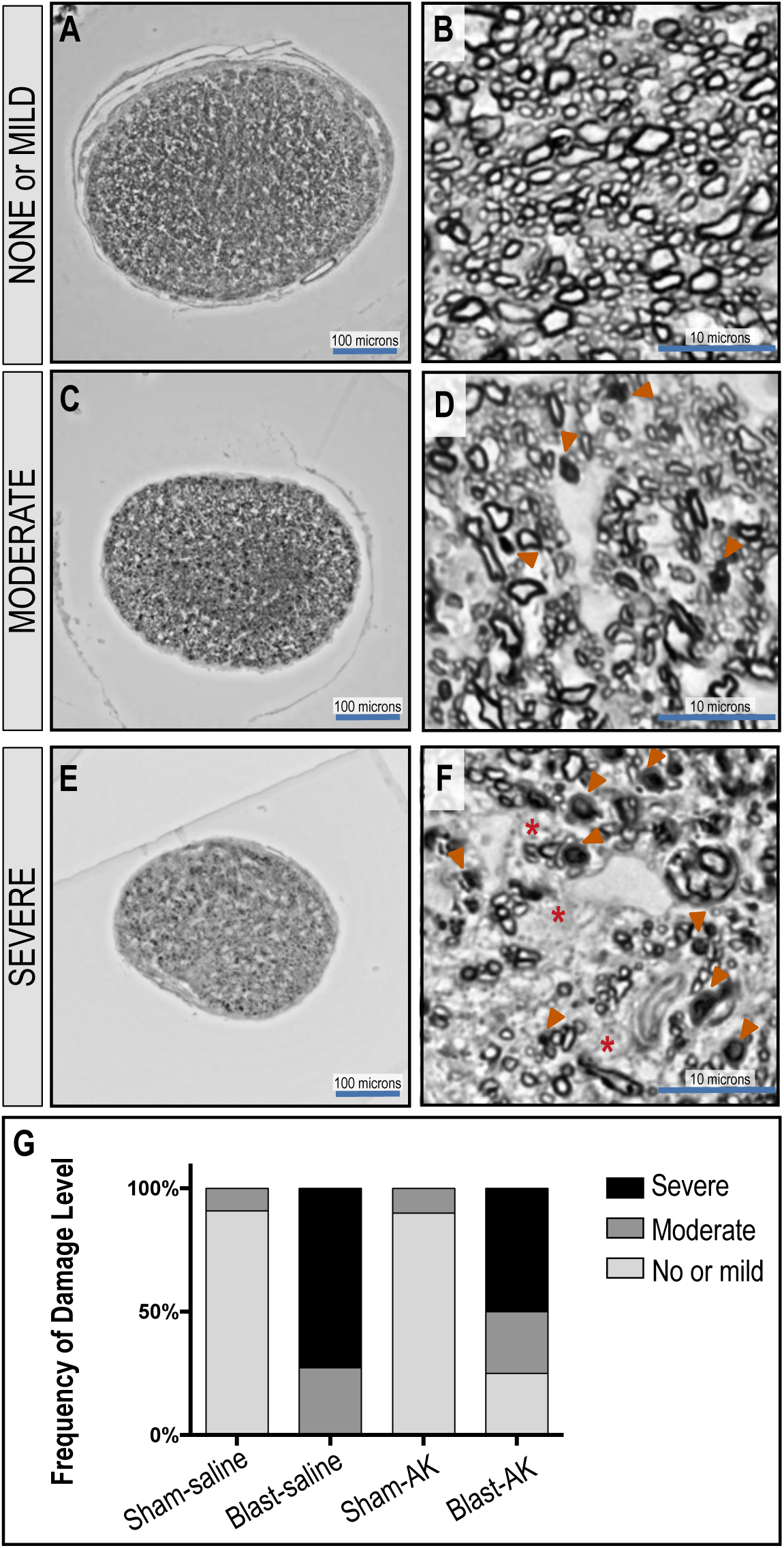
Anakinra treatment supports survival of axon bundles in the optic nerve after blast. The degree of neurodegeneration was assessed by analysis of the optic nerve axonal damage across sections. Representative low and high magnification image of ipsilateral optic nerve cross sections. Damage levels based on grade. Grade 1: healthy-mild damage (**A**,**B**), Grade 2: moderate damage (**C, D**), Grade 3: severe damage (**E**,**F**). Arrow heads indicate PPD-positive and infilled dead/dying axons; asterics indicate glal scar formation adjacent to glial cell nuclei. Frequency of damage level demonstrates that treatment with anakinra prevents damage of the optic nerve (**G**).

PERG amplitudes were utilized to measure the functional signaling capacity of RGCs. The prerecorded baseline peak to trough amplitude for the ipsilateral eyes in the sham-saline, blast-saline, sham-anakinra, and blast-anakinra groups were 22.88 ± 1.055 µV, 22.51 ± 1.429 µV, 23.65 ± 1.506 µV, and 20.16 ± 1.007 µV, respectively, with no significant differences found when comparing groups to the sham-saline as the control (**Fig 11A**). 4 weeks after injury, only the blast-saline group had significantly decreased PERG peak to trough amplitudes (*P*=0.0170) when compared to the sham-saline group, suggesting that the blast group treated with anakinra had partially rescued RGC signaling (SS=19.93 ± 1.546 µV; BS=13.44 ± 1.768 µV; SA=22.62 ± 1.74 µV; BA=16.22 ± 1.37 µV; **Fig 11B**).

**Figure 11.**
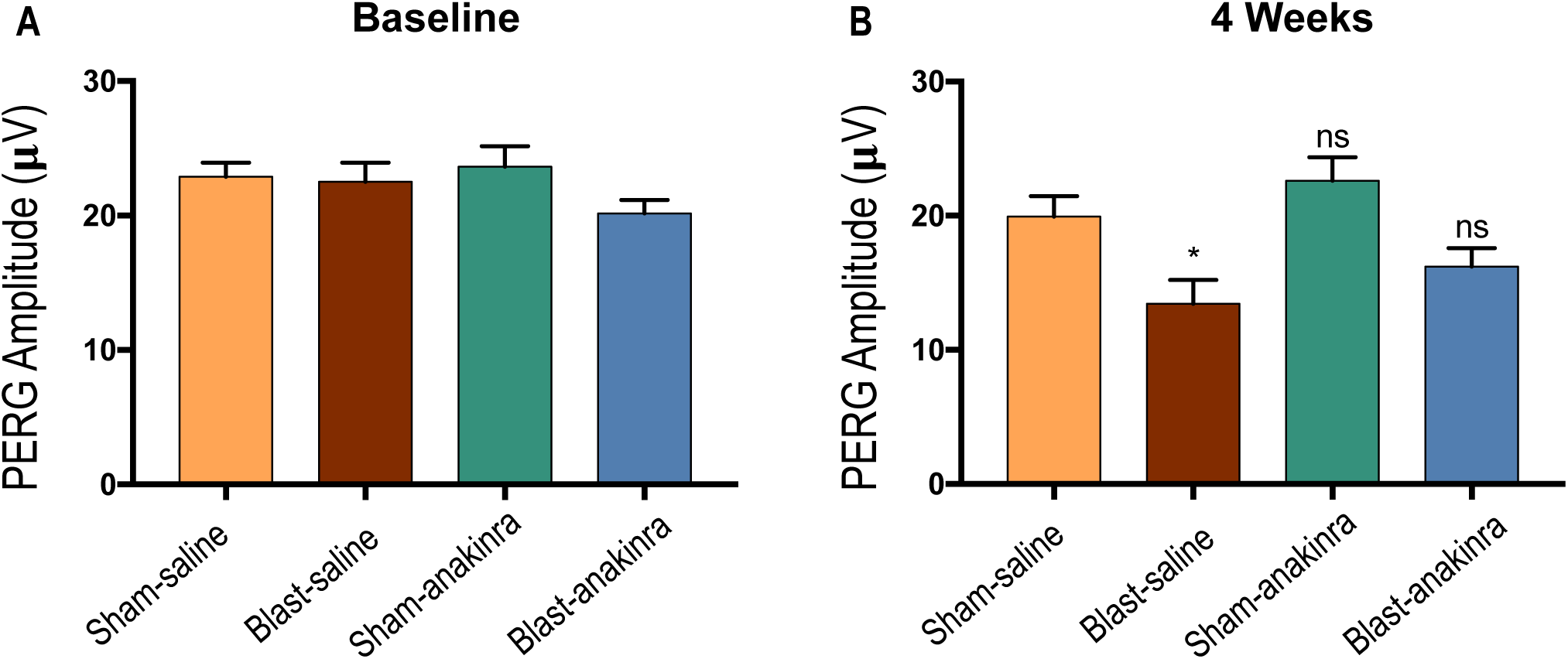
Blast-induced functional RGC damage post repeated bTBI is partially rescued by anakinra. Significant difference compared to sham-saline group using Kruskal-Wallis test with Dunn’s posttest. \**P* <0.05. (RGC: retinal ganglion cell; PERG: pattern-evoked electroretinography).

To evaluate the structural integrity of the retina, we used SD-OCT to measure the *in vivo* thickness of the RGC complex (RGC cell bodies, axons, and dendtrites). The prerecorded baseline RGC complex thickness for the ipsilateral eyes in the sham-saline, blast-saline, sham-anakinra, and blast-anakinra groups were 76.07 ± 0.6511 µm, 75.58 ± 0.7159 µm, 75.01 ± 0.8511 µm, and 73.83 ± 0.6522 µm, respectively, with no significant differences found between groups (**Fig 12A**). Four weeks after injury the sham-saline and sham-anakinra groups remained statistically unchanged from their baseline values, with average changes in RGC complex thickness of −0.4734 ± 0.532 µm and −0.6475 ± 0.9451 µm, respectively (representative images **Fig 12C and E**). By contrast, blast injury elicited a decrease in RGC complex thickness in the blast-saline group (**Fig 12D**) by -10.61 ± 1.131 µm that was significantly different from all other groups (P<0.0001, **Fig 12B**). After treatment with anakinra, the blast-anakinra group exhibited preservation of the RGC complex, with a change from baseline of −3.1714 ± 1.045 µm (**Fig 12F**), suggesting that anakinra helped to maintain the structural integrity of the RGCs.

**Figure 12.**
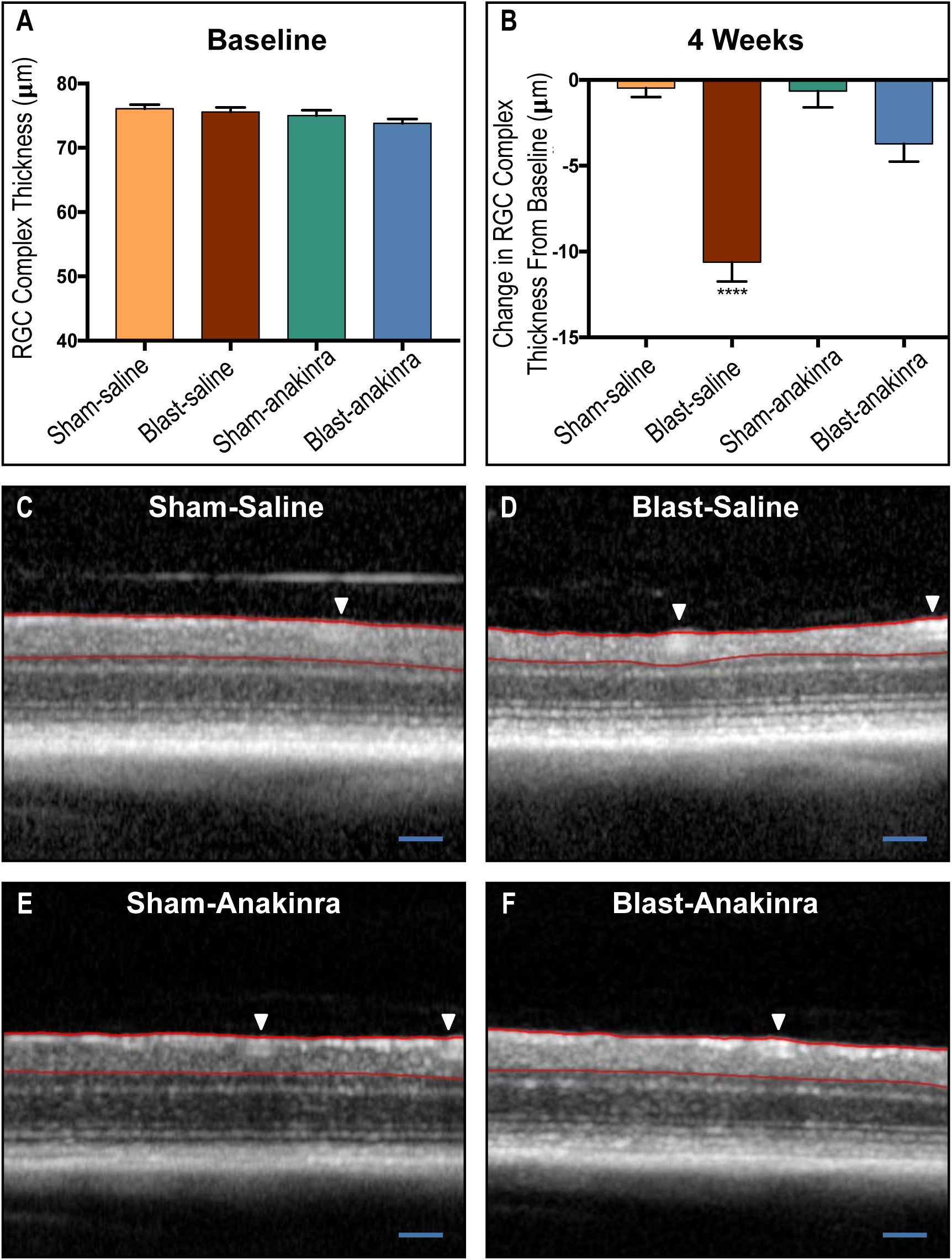
RGC complex loss due to bTBI is partially rescued by anakinra. The area between the red lines is the RGC complex thickness measured. No significant differences between groups were seen at baseline (**A)**. At 4 weeks post injury, both sham groups and the blast-anakinra group had significantly less of a change in RGC complex from baseline when compared to the blast-saline group, with no significant differences found between other groups **(B)**. Representative OCT images of sham-saline **(C)**, blast-saline **(D)**, sham-anakinra **(E)**, and blast-anakinra **(F)**. Arrows indicate blood vessels exclude from analysis. All data from ipsilateral retinas. One-way ANOVA with Dunnett’s posttest comparing all means (\*\*\*\**P*<0.0001). Scale bar: 200 µm.

## Discussion

We have shown that repeated bTBI induces acute retinal inflammation, glial cell activation, and chronic neuronal dysfunction, which can be mitigated in part by IL-1 pathway blockade via anakinra. The acute inflammation we see in the retina and brain is seen in several other mouse models and human TBI (Hutchinson, O’Connell et al. 2007, Simon, McGeachy et al. 2017, Newell, Todd et al. 2018), highlighting the pervasiveness of damaging secondary inflammation across models.

We previously identified the presence of inflammatory cells histologically in the retina after blast injury (Evans, Newell et al. 2018), and in this study found activation of microglia and Müller glia in response to injury. As the retina is an immunopriviledged site, this hyperactivation of resident microglia (and possibly peripheral leukocytes due to blood-retinal-barrier disruption after blast) (Hue, Cho et al. 2016) may significantly contribute to retinal tissue damage without an efficient way to regulate this immunological cascade. Microglia can be activated by RGC damage, resulting in proliferation and migration to areas of damage within the retina (Cuenca, Fernandez-Sanchez et al. 2014). The morphological activation and increased number of microglia that we observed near the RNFL in our model suggests that microglia are responding to blast-induced RGC damage and that their activation is further contributing to neurodegeneration and cell death.

In the healthy retina, Müller glia prevent photoreceptor and neuronal cell death via the secretion of neurotrophic factors, growth factors, and cytokines (Bringmann, Pannicke et al. 2006, Bringmann, Iandiev et al. 2009). After blast injury, Müller glia demonstrated increased distribution of GFAP immunoreactivity, indicative of an injury response and retinal stress. Both activated microglia and Müller glial can contribute to retinal damage, as they are activated by and can continue to propogate damaging inflammatory signaling, acting as immunocompentent cells. Modulation of glial activation through IL-1 blockade could be responsible for the improved visual outcomes in the setting of anakinra treatment after blast trauma. Histologic analysis revealed that anakinra reduced the overall cellular activation of microglia and Müller glia 4 weeks after injury. This treatment maintained the signaling capability as well as the structural integrity of the RGCs and optic nerve. Preservation of RGC function and structure is necessary to maintain visual transduction from the retina to higher visual processing centers in the brain.

This study demonstrated that damage to retinal ganglion cells can be detected after blast injury using noninvasive functional and structural tests, and that anti-IL-1 therapy through anakinra can confer partial protection to these cells. Pharmacologic blockade of retinal inflammation could potentially be utilized to improve visual outcomes and quality of life in bTBI patients. Anakinra has an excellent safety profile even when used in individuals with active bacterial infections making it an ideal cytokine blocker in this scenario as blast injuries can be accompanied by penetrating trauma depending on the device used and proximity to the blast epicenter. Future studies will be aimed towards dissecting the IL-1 pathway to gain a greater understanding of each molecule in order to therapeutically manipulate them after TBI.

## Acknowledgements

AGB and VBM are supported by the NIH grant EY26682. PJF is funded by the NIH/NIAMS R01AR059703 and R01NS098590 NIH/NINDS.

MMH is supported by the Department of Veterans Affairs grant RX000952, Department of Defense grant W81XWH-14-1-0583, and the Iowa City Center for the Prevention and Treatment of Visual Loss. AHB is supported by training grant T32 DK112751-01. Additional support was provided by NIH/NEI Center Support Grant to the University of Iowa (P30 EY025580). The contents of this manuscript do not represent the views of the U.S. Department of Veterans Affairs, Department of Defense, or the U.S. Government.

The authors declare no competing financial interests.

